# Methylation reprogramming associated with aggressive prostate cancer and ancestral disparities

**DOI:** 10.1101/2025.04.16.649098

**Authors:** Jenna Craddock, Pavlo Lutsik, Pamela X.Y. Soh, Melanie Louw, Md. Mehedi Hasan, Sean M. Patrick, Shingai B.A. Mutambirwa, Phillip D. Stricker, Hagen E.A. Förtsch, HEROIC PCaPH Africa1K, M.S. Riana Bornman, Clarissa Gerhäuser, Vanessa M. Hayes

## Abstract

African men are disproportionately impacted by aggressive prostate cancer (PCa). Key to this disparity both genetic and environmental factors, alluding to epigenetic modifications. However, African-inclusive prostate tumour DNA methylation studies are lacking. Assembling a multi-geo-ancestral prostate tissue cohort, including men with (57 African, 48 European, 23 Asian) or without (65 African) PCa, we interrogate for genome-wide differential methylation. Overall, methylation appears to be driven by ancestry over geography (152 southern Africa, 41 Australia). African tumours show substantial heterogeneity, with universal hypermethylation indicating epigenetic silencing, encompassing PCa suppressor genes and enhancer-targeted binding motifs. Conversely, African tumour-associated heterochromatic hypomethylation suggests permissive chromatin remodelling, with developmental pathway activation via enhancer targets. Taken together, we show methylation aberrations favour metastatic growth, genomic instability and disease aggressiveness in African tumours, which we hypothesise is driven by extensive plasticity of intergenic regulatory regions.

## Introduction

Prostate cancer (PCa) is a significant global health concern. The second most frequently diagnosed cancer among men worldwide, PCa was responsible for approximately 1.47 million new cases and 397,000 deaths in 2022^1^. However, PCa disproportionately burdens men of African ancestry and/or from Africa. A focus on the United States has revealed that African American men display a 2- and 4.3-fold increased risk for PCa mortality compared to their European and Asian counterparts, respectively^2^. Conversely, Sub-Saharan Africa has received the top four of five ranks for global PCa mortality rates, with southern Africa leading the charge at a 4-fold increase^1^. This disparity underscores the urgent need for research efforts tailored to the African context. Yet, Sub-Saharan African populations remain underrepresented in PCa research^3^, with limited focus on exomic^4^ and whole genome interrogation^5,6^, the role of epigenetics remains unexplored.

DNA methylation aberrations are some of the earliest, most stable and frequent molecular changes to occur in PCa^7^. More frequent than genetic alterations, they are preserved throughout progression and metastatic development. With no known significant PCa modifiable risk factors^8^, somatic DNA methylation profiling holds potential to identify contributing environmental carcinogens^9^. Notable DNA methylation differences in prostate tumours between ancestries include *TIMP3* hypermethylation in Black over White Americans^10^, while glaring inconsistencies include prostate tumour *CD44* hypermethylation^11^ *versus* no differential methylation^12^, and *PMEPA1* hypermethylated^10^ *versus* hypomethylated^13^ for Black Americans. Well-established for being epigenetically silenced in PCa, *GSTP1* appears to be universally hypermethylated across prostate tumours regardless of patient ancestry^14^. While methylation profiles distinguishing prostate tumour from benign tissue appear to be similar across American ancestries^15^, dysregulation of the androgen receptor signalling pathway coupled with decreased expression of immune-related genes has been reported to be African-derived tumour specific^16^. Inconsistency between studies likely arise from small sample sizes, varying methylation profiling methods, non-biological factors or the heterogenous nature of populations investigated^17^, including historical genetic admixture^18^. Data for Sub-Saharan Africa appears to be lacking.

Additionally, European bias associated with DNA methylation profiling array design, including the Illumina MethylationEPIC BeadChips, raises concerns with regards to poor CpG probe hybridization within genetically highly diverse African populations^19^. Recognizing this caveat, Zhang et al.^20^ recently characterized Illumina probes in global populations using the 1000 Genomes Project data. Including East and West African, and predominantly West African ancestral Black American and African Caribbean populations^18^, southern Africans at greatest risk for PCa mortality^1^ and representing the greatest regional genetic diversity^21^, are yet to be represented.

In this study of prostate tissue derived from a unique cohort of 193 individuals biased towards southern Africans (122 African, 30 European), with further comparison to Australians (18 European, 23 Asian), we generate not only a southern African-relevant methylome-wide EPIC array filtering resource, but also interrogate for both geo-ancestral and tissue-specific differential DNA methylation. We find prostate tumour methylation to be driven by patient ancestry over geography, while African tumours show significant hypermethylation, heterogeneity and elevated gene silencing. The identification of African-specific PCa gene targets, as well as greater African-derived tumour immune function impairment, provides further rationale for the importance of African inclusion in epigenetic studies focused on reducing the global PCa burden.

## Results

### African-inclusive DNA methylation array limitations, confounders and cohort characterisation

Avoiding unreliable hybridization with consequential DNA methylation mismeasurement^19^, single nucleotide polymorphism (SNP)-overlapping probe filtering is a common data processing feature in current workflows. Established using largely European-derived resources, although a more recent African-inclusive strategy has been proposed^20^, our focus on genetically diverse southern Africans raises concerns. Using published whole genome sequenced blood-derived data (average coverage 43X; range 32-69X) from 99 genetically confirmed southern Africans, with germline single nucleotide variants (SNVs) and insertions/deletions <50 bases (indels) called referencing GRCh38^5^, we generate a southern African-relevant variant filtering resource (minor allele frequency (MAF) >0.01), comprising 56,280 SNVs and 3,623 indels (EPICv1, Supplementary Data 1-3), and 50,786 SNVs and 3,087 indels (EPICv2, Supplementary Data 4-6). Our workflow is outlined in Supplementary Fig. 1.

Using our SNP-filtering resource, we further evaluate for African-specific discrepancies in probe content between EPIC array versions. Observing a greater proportional loss for EPICv1 (6.91% or 59,903) over EPICv2 sites (5.75% or 53,873), SNP overlap was identified in three categories (MAF > 0.01): (i) target CpG sites (45,669 EPICv1; 33,772 EPICv2), (ii) single base extension (SBE) sites of Type I probes (2,359; 746), and (iii) overlapping the probe body within 5bp of the target CpG (22,103; 25,029). Our African EPICv1 assessment reveals a 1.4-fold overlap with target CpG sites, and 1.6-fold with SBE sites when compared with equivalent European-relevant confounding^22^. Due to African variant confounding and compared with EPICv1, EPICv2 displays greater proportional loss within gene bodies (49.50% EPICv1; 59.14% EPICv2) (Fig. 1a), non-CpG island (CGI) regions (72.97%; 79.08%) (Fig. 1b) and across Type II probes (84.23%; 89.49%) (Fig. 1c) and FANTOM5 enhancers (3.80%; 3.98%) (Fig. 1d). This observation is likely explained by elevated coverage over said regions and greater Type II probe content (abundant over Type I across platforms) on EPICv2 over EPICv1^23^. Within gene regions, although probe distribution across platforms is dominant in transcription start sites (TSSs)^23^, probe loss does not reflect this distribution – a positive finding.

**Fig. 1:**
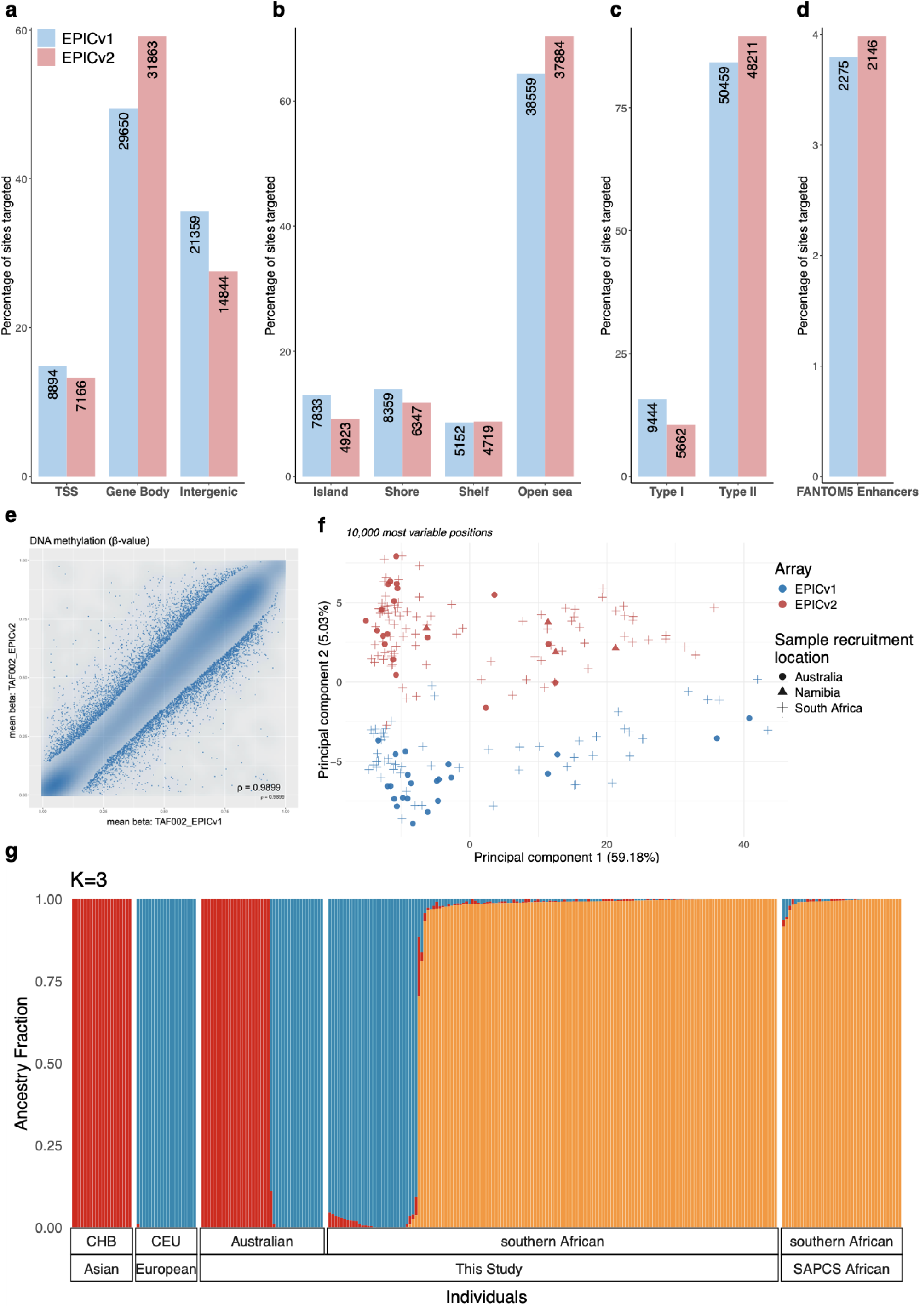
DNA methylation geo-ancestral cohort analysis using the EPICv1 and EPICv2 arrays. **a–d** Percentage (bars) and number (labels) of probe sites, distributed on EPICv1 *versus* EPICv2, identified to be confounded by African genomic diversity relative to **a** gene regions, **b** CpG island context, **c** CpG probe type and **d** overlapping FANTOM5 enhancers. Percentages are calculated relative to total African-confounding probe loss per platform. **e** Scatterplot showing the correlation between methylation measurements from EPICv1 and EPICv2 for African replicate pair TAF002. **f** Principal component analysis (PCA) of a 193-sample merged and normalized EPICv1/EPICv2 dataset across the 10,000 most variable positions. Samples are annotated according to array version: EPICv1 (*n* = 78) and EPICv2 (*n* = 115); and the location from which samples were recruited: Australia (*n* = 41), Namibia (*n* = 4) and South Africa (*n* = 148). **g** Admixture plot (*K* = 3, cross-validation error = 0.524) replicated in 10/10 runs, including 40 southern Africans from the Southern African Prostate Cancer Study (SAPCS), both including and excluding for Khoe-San fractions^5^, and 20 Europeans (CEU) and 20 Chinese (CHB) from the Human Genome Diversity Project (HGDP) and 1000 Genomes Project (1KGP) subset of gnomAD v3.12^24^, together with our geo-ancestral cohort (*n* = 192, with exclusion of a single African with insufficient genome sequencing coverage).

To evaluate cross-platform reproducibility, we profiled a subset of matched African sample pairs on both the EPICv1 and EPICv2 array (*n* = 7), observing a high β-value correlation between the EPIC arrays (all ρ >0.9838) (Fig. 1e, Supplementary Fig. 2). Suggesting African-specific reproducibility and suitability for cross-platform integration, it was surprising that our 193-sample merged dataset (78 EPICv1; 115 EPICv2) (Table 1) showed, after normalization and background subtraction, significant array version confounding with principal component analysis (PCA) (Fig. 1f, Supplementary Table 1). Notably, this separation cannot adequately be explained by possible variation across different sample collection sites. Using our genetically confirmed through ancestral fraction analyses (122 African, 48 European, 23 Asian; Fig. 1g) and geographically (148 South Africa, 4 Namibia, 41 Australia) diverse cohort to eliminate this confounding contribution, we approached our analyses using a larger discovery *versus* smaller validation cohort derived from alternate array types (Table 1). As such, our “ancestry-associated” (African *versus* non-African) analysis included an EPICv1 discovery (*n* = 70) and EPICv2 validation cohort (*n* = 57), while our “tumour-associated” (tumour *versus* normal/benign tissue) analysis included an EPICv2 discovery (*n* = 93) and EPICv1 validation cohort (*n* = 29). We present only discovery cohort observations that were consistent with the validation cohort.

**Table 1.**
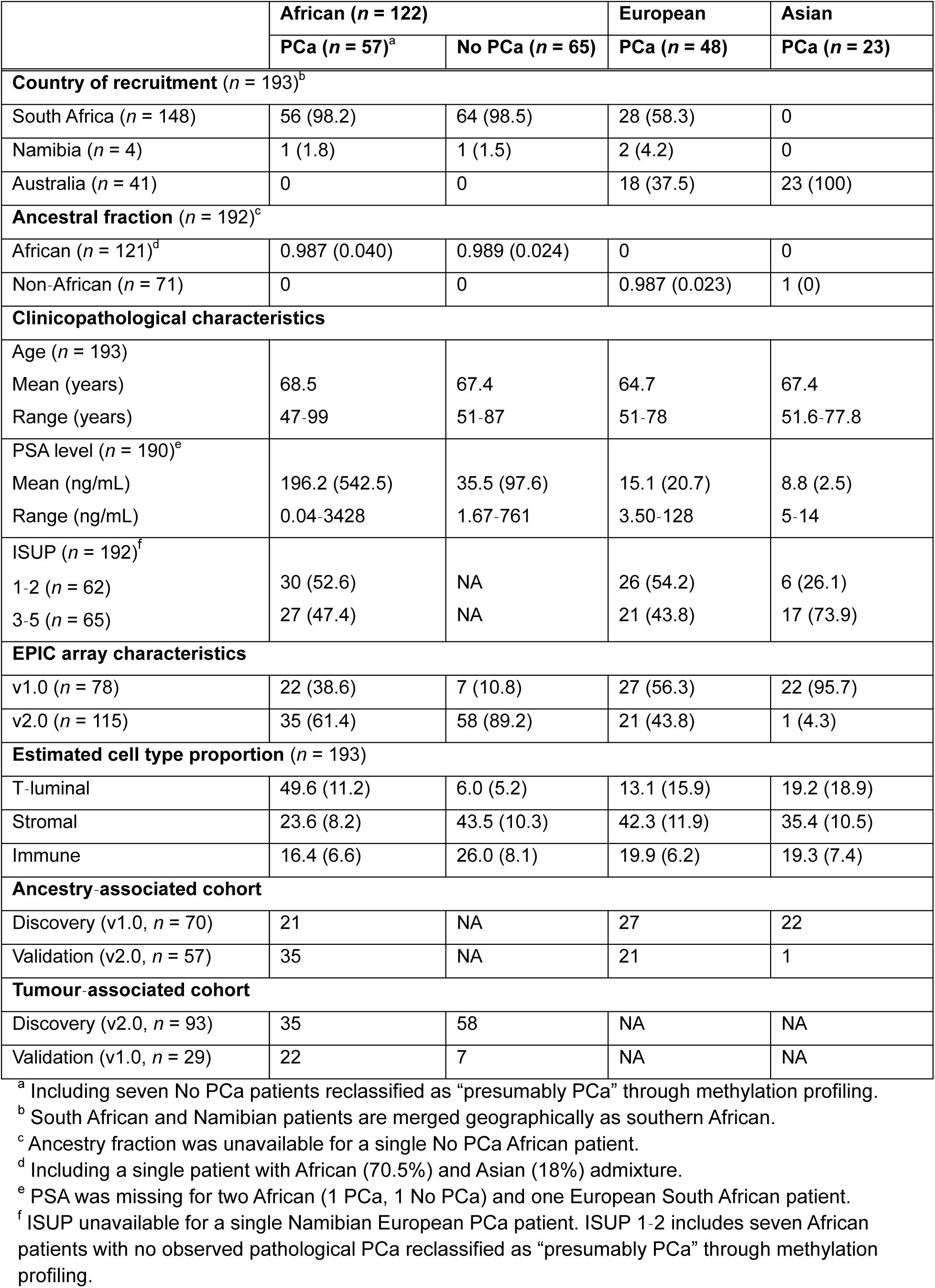
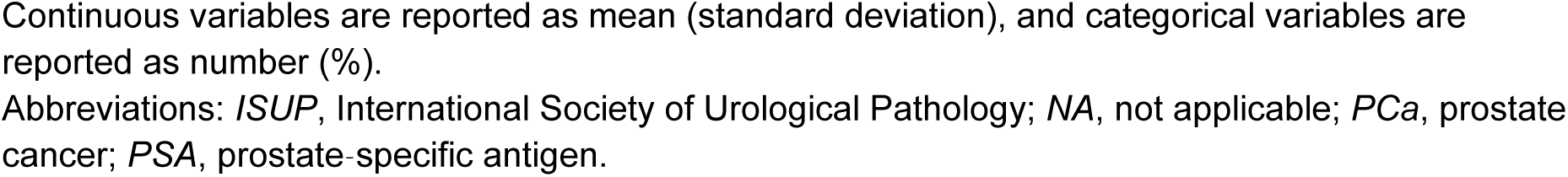
Study participant geo-ancestries and clinicopathological prostate cancer characteristics, including tissue associated EPIC array-based features.

|  | African (*n* = 122) |  | European | Asian |
| --- | --- | --- | --- | --- |
|  | PCa (*n* = 57)<sup>a</sup> | No PCa (*n* = 65) | PCa (*n* = 48) | PCa (*n* = 23) |
| **Country of recruitment (*n* = 193)<sup>b</sup>** |  |  |  |  |
| South Africa (*n* = 148) | 56 (98.2) | 64 (98.5) | 28 (58.3) | 0 |
| Namibia (*n* = 4) | 1 (1.8) | 1 (1.5) | 2 (4.2) | 0 |
| Australia (*n* = 41) | 0 | 0 | 18 (37.5) | 23 (100) |
| **Ancestral fraction (*n* = 192)<sup>c</sup>** |  |  |  |  |
| African (*n* = 121)<sup>d</sup> | 0.987 (0.040) | 0.989 (0.024) | 0 | 0 |
| Non-African (*n* = 71) | 0 | 0 | 0.987 (0.023) | 1 (0) |
| **Clinicopathological characteristics** |  |  |  |  |
| Age (*n* = 193) |  |  |  |  |
| Mean (years) | 68.5 | 67.4 | 64.7 | 67.4 |
| Range (years) | 47-99 | 51-87 | 51-78 | 51.6-77.8 |
| PSA level (*n* = 190)<sup>e</sup> |  |  |  |  |
| Mean (ng/mL) | 196.2 (542.5) | 35.5 (97.6) | 15.1 (20.7) | 8.8 (2.5) |
| Range (ng/mL) | 0.04-3428 | 1.67-761 | 3.50-128 | 5-14 |
| ISUP (*n* = 192)<sup>f</sup> |  |  |  |  |
| 1-2 (*n* = 62) | 30 (52.6) | NA | 26 (54.2) | 6 (26.1) |
| 3-5 (*n* = 65) | 27 (47.4) | NA | 21 (43.8) | 17 (73.9) |
| **EPIC array characteristics** |  |  |  |  |
| v1.0 (*n* = 78) | 22 (38.6) | 7 (10.8) | 27 (56.3) | 22 (95.7) |
| v2.0 (*n* = 115) | 35 (61.4) | 58 (89.2) | 21 (43.8) | 1 (4.3) |
| **Estimated cell type proportion (*n* = 193)** |  |  |  |  |
| T-luminal | 49.6 (11.2) | 6.0 (5.2) | 13.1 (15.9) | 19.2 (18.9) |
| Stromal | 23.6 (8.2) | 43.5 (10.3) | 42.3 (11.9) | 35.4 (10.5) |
| Immune | 16.4 (6.6) | 26.0 (8.1) | 19.9 (6.2) | 19.3 (7.4) |
| **Ancestry-associated cohort** |  |  |  |  |
| Discovery (v1.0, *n* = 70) | 21 | NA | 27 | 22 |
| Validation (v2.0, *n* = 57) | 35 | NA | 21 | 1 |
| **Tumour-associated cohort** |  |  |  |  |
| Discovery (v2.0, *n* = 93) | 35 | 58 | NA | NA |
| Validation (v1.0, *n* = 29) | 22 | 7 | NA | NA |

<sup>a</sup> Including seven No PCa patients reclassified as “presumably PCa” through methylation profiling.
<sup>b</sup> South African and Namibian patients are merged geographically as southern African.
<sup>c</sup> Ancestry fraction was unavailable for a single No PCa African patient.
<sup>d</sup> Including a single patient with African (70.5%) and Asian (18%) admixture.
<sup>e</sup> PSA was missing for two African (1 PCa, 1 No PCa) and one European South African patient.
<sup>f</sup> ISUP unavailable for a single Namibian European PCa patient. ISUP 1-2 includes seven African patients with no observed pathological PCa reclassified as “presumably PCa” through methylation profiling.
Continuous variables are reported as mean (standard deviation), and categorical variables are reported as number (%). Abbreviations: *ISUP*, International Society of Urological Pathology; *NA*, not applicable; *PCa*, prostate cancer; *PSA*, prostate-specific antigen.

Overall, we observed vast DNA methylation heterogeneity amongst our African samples. Notably, seven African samples histologically classified as “non-tumour” revealed DNA methylation profiles resembling tumour samples (Supplementary Fig. 3). Speculating that cancerous tissue may have been missed during routine biopsy, these samples were recharacterized as “presumably PCa” in downstream analyses. Additionally, as we observed β-value noise from irrelevant cell types in our African samples, only those belonging to the highest tumour purity quartile were included (reducing our initial quality-controlled study cohort from 282 to 193, see Methods). Consequently, ancestry is significantly confounded with certain cell type proportions, necessitating validation of ancestry-associated differential methylation findings, as described above.

### Ancestry-associated prostate tumour differential methylation discovery

EPICv1 “ancestry-associated discovery cohort” (Table 1), was used to perform geo-ancestral differential methylation analyses between prostate tumours derived from patients of African (*n* = 21, all southern African) and non-African ancestry (*n* = 49; 27 European southern Africans, 22 Asian Australians). Initial PCA visualization revealed that despite shared geography between Black and White southern Africans, tumour-driven DNA methylation grouping appears to be largely driven by ancestry, with PC1 explaining 78.76% of the variance in the methylation data and most strongly associating with ancestry, following tumour purity, unsurprising given exclusively high African tumour purity inclusion (Fig. 2a, Supplementary Fig. 4, Supplementary Table 2). While little difference was observed between European and Asian tumours, African tumours showed great between-sample heterogeneity not adequately explained by tumour content (Supplementary Fig. 5). Notably, no significant differences in global methylation at repetitive elements Alu, LINE-1 and LTR were observed (Supplementary Fig. 6).

**Fig. 2:**
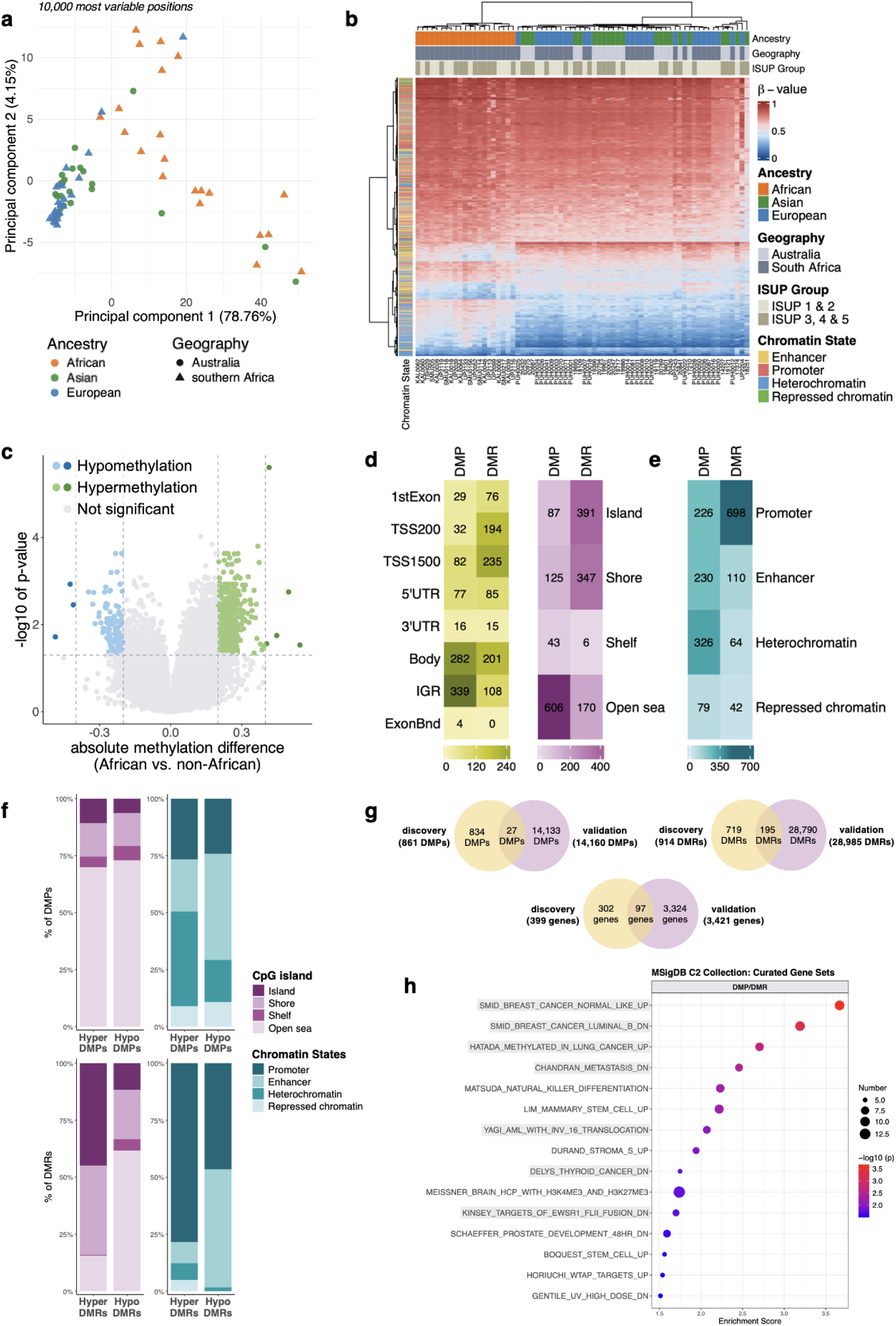
Differential DNA methylation analysis for African-*versus* non-African-derived prostate tumours. **a** PCA for the 10,000 most variable positions for the EPICv1 “ancestry-associated discovery cohort” by ancestry: African (*n* = 21), European (*n* = 27) and Asian (*n* = 22), and geography: South Africa (*n* = 48) and Australia (*n* = 22). **b** DMP cluster analysis heatmap by ancestry, geography, ISUP grade group and chromatin state. Rows represent CpG sites (*n* = 861) and columns, patients (*n* = 70). The methylation level is represented by a β-value between 0 (completely unmethylated, blue) and 1 (fully methylated, red). **c** Volcano plot representing 861 DMPs by ancestral (African *versus* non-African) significance (BH FDR *p* <0.05, |Δβ| ≥20% and 40%, dashed lines), including hypermethylated (green) or hypomethylated (blue), compared with non-significant (grey) DMPs. **d** Heatmap showing gene region (green) and CpG island-related region (purple) enrichment for DMP- and DMR-related CpG sites across various contexts. **e** Heatmap showing chromatin state enrichment of DMP- and DMR-related CpG sites across various contexts. **f** Stacked bar graphs of the percent overlap of DMPs and DMRs with various CpG island-related (purple) and chromatin state (blue) contexts. **g** Venn diagrams illustrating significant DMP, DMR and annotated gene agreement between the ancestry-associated discovery and validation cohorts. **h** MSigDB (C2) enrichment for validated genes collectively annotated to hypermethylated DMPs and DMRs for the top 15 terms (minimum 5 genes), with cancer-related terms highlighted in grey. Point size reflects the number of genes associated with each gene set. Association strength is denoted by –log_10_(*p*-value). Abbreviations: *DMP*, differentially methylated position; *DMR,* differentially methylated region; *ExonBnd*, within 20 bases of an exon boundary; *IGR*, intergenic region; *ISUP*, International Society of Urological Pathology; *MSigDB*, Molecular Signatures Database; *PCA*, principal components analysis; *TSS*, transcription start site, *UTR* untranslated region; *|Δβ|*, absolute difference in mean methylation.

Overall, we identified 861 differentially methylated positions (DMPs) associated with patient ancestry (African *versus* non-African BH false discovery rate (FDR) *p* <0.05, |Δβ| ≥20%) (Supplementary Table 3), with the top 10 impacting cancer associated genes (Table 2). Further cluster analysis (861 DMPs) reiterates ancestral over geographic groupings and extensive African-specific tumour heterogeneity (Fig. 2b). Compared to non-Africans, 721 DMPs (83.74%) are hypermethylated and 140 DMPs hypomethylated in Africans (Fig. 2c). Although gene expression regulation by DNA methylation is often discussed in the context of protein-coding relevant regions like CGIs and neighbouring areas (within 2kb i.e. CGI shore), as well as promoter regions, it is also true that enhancer elements are enriched with disease-associated epigenetic changes^25^. Notably, 39.37% of the ancestry-associated DMPs were located in intergenic regions (IGR), coupled with predominant distribution in open sea regions (70.38%) (Fig. 2d). Using a prostate tumour-derived chromatin state annotation^26^, we found the ancestry-related DMPs to be primarily enriched in heterochromatin (37.86%, *p* = 7.27 × 10^−5^, Fisher’s exact test), followed by enhancers (26.71%), including active and primed non-prostate-(8.48%, *p* = 2.26 × 10^−3^ and 3.60%, *p* = 1.52 × 10^−11^, respectively) and prostate lineage-specific (7.08%, *p* = 3.58 × 10^−9^ and 4.53%, *p* = 6.99 × 10^−5^, respectively) enhancers, as well as in promoters (26.25%), particularly active non-prostate lineage promoters (11.38%, *p* = 4.29 × 10^−5^), and repressed chromatin (9.18%, *p* = 0.03) (Fig. 2e, Supplementary Table 3). While hypermethylated DMPs are primarily located in heterochromatin (41.61%, *p* = 3.95 × 10^−7^) and hypomethylated DMPs, in enhancers collectively (46.43%, *p* = 6.27 × 10^−4^), promoters and repressed chromatin show near equal hyper- and hypomethylation distribution (promoter 26.63% and 24.29%, repressed chromatin 8.88% and 10.71%, respectively) (Fig. 2f, Supplementary Fig. 7).

**Table 2.**
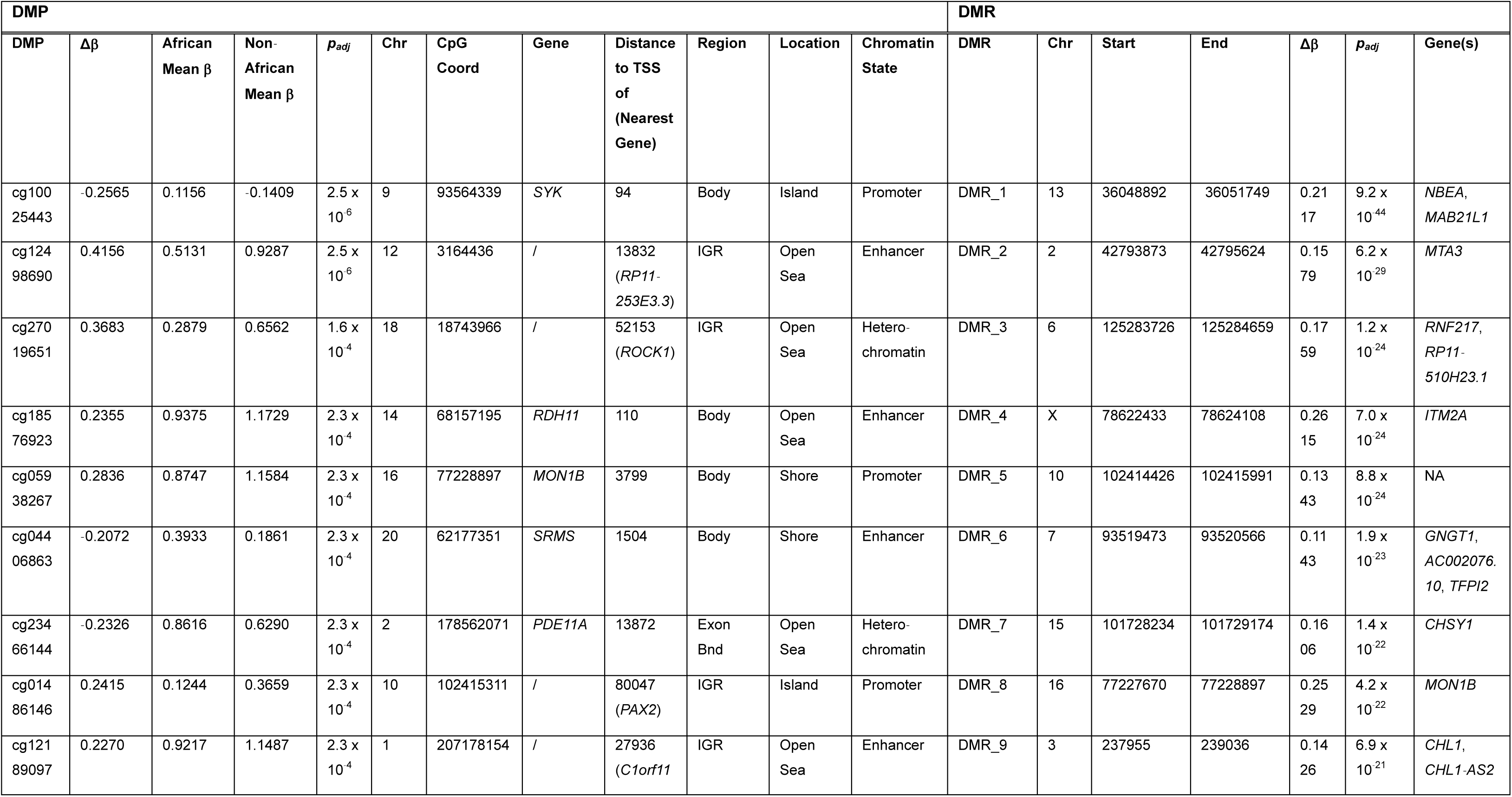
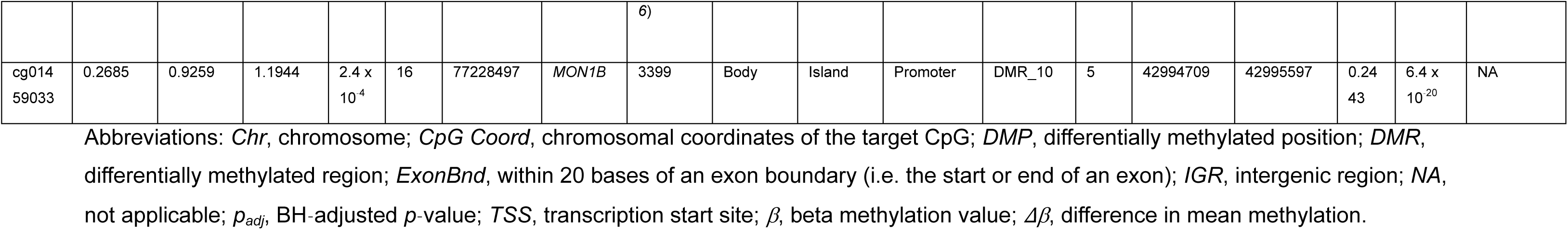
Top 10 significant differentially methylated positions (DMPs) and regions (DMRs) identified between African and non-African prostate tumours.

Using DMRcate^27^, we further identified 110 differentially methylated regions (DMRs) comprising 914 CpGs (FDR *p* <0.05, |Δβ| ≥10%), with the vast majority (91.82%) hypermethylated in African over non-African tumours (Supplementary Tables 4-5). Again, all top 10 significant DMRs impact genes having oncogenic potential (Table 2), with DMRs largely overlapping promoter regions and CGIs (Fig. 2d-e). However, while most hypermethylated DMRs are located in CGIs and shores (44.96% and 39.11%, respectively), hypomethylated DMRs are more common in non-CGI regions (61.67%) (Fig. 2f), although hypomethylation overlap with common partially methylated domains (PMDs) was minimal (8.33%). Relative to annotated chromatin states, hypermethylated (78.45%, *p* <2.2 × 10^−16^) over hypomethylated (46.67%, *p* = 0.10) DMRs are more common to promoters, with hypomethylated DMRs showing predominant enhancer distribution (51.67%, *p* = 6.78 × 10^−3^). More specifically, hypomethylated DMRs are found within active non-prostate-(20%) and prostate lineage-specific (18.33%) promoters and primed non-prostate-(15%) and prostate lineage-specific enhancers (16.67%) (Supplementary Fig. 7).

Exclusively considering significant differential methylation overlap with our ancestry-associated validation cohort (Fig. 2g) for functional enrichment analysis, and further filtering to include genes that display a correlation between methylation and expression in the prostate (see Methods), we examined whether DMPs and DMRs were collectively enriched in biological pathways of interest presented in curated gene sets from the Molecular Signatures Database (MSigDB)^28^. Within the MSigDB C2 collection, hypermethylated DMPs/DMRs show enrichment for cancer-related gene sets including genes reported as: downregulated in metastatic PCa and in breast cancer, hypermethylated in lung cancer (Fig. 2h), and possessing histone deacetylase (HDAC) activity (Supplementary Table 6). Additionally, hypermethylated DMPs/DMRs were enriched for several oncogenic (C6 collection) and immunologic signature (C7) gene sets. Although poorly enriched, hypomethylated DMPs were likewise enriched in immune- and cancer-related gene sets, with notable inclusion of genes upregulated in breast cancer and in mammary stem cells. Additionally, hallmark gene sets include genes involved in epithelial to mesenchymal transition (EMT). Notably, few pathways were significant after controlling for a 5% FDR. Using data from the GeneHancer ‘Double Elite’ list^29^, we further verified DMP and DMR overlap with 33 validated promoter/enhancer elements. Two or more sources of evidence suggest a high likelihood of interaction between these regions and 63 target genes collectively (Supplementary Table 7). Target genes of differentially methylated enhancers were notably enriched in developmental pathways, various gene sets of known transcription factor (TF) targets, as well as in (prostate) cancer-associated pathways (Supplementary Table 8).

Validated DMP and DMR CpGs were further annotated to 97 genes (Fig. 2g), of which 16 were both DMP- and DMR-related. Besides 12 known PCa-associated genes (Supplementary Table 9), we found four genes to be of particular interest as potentially unknown African-specific PCa targets, namely *GALM*, *EVC2*, *CHSY1* and *SPDYA* (Fig. 3). Of the genes reported to be frequently differentially methylated between African American and European American prostate tumours (e.g. *RARB*, *TIMP3*)^17^, none were significantly differentially methylated in our cohort.

**Fig. 3:**
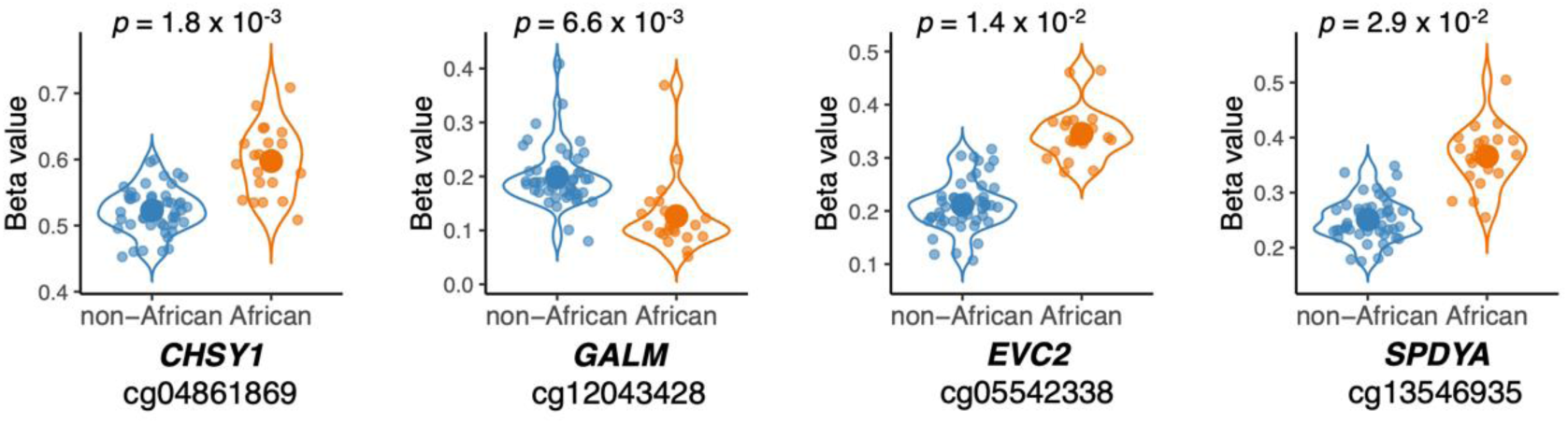
African *versus* non-African CpG DNA methylation plots for four individual potentially unknown African-specific prostate cancer (PCa) targets. Noteworthy significant differentially methylated positions (DMPs) between African (orange) and non-African (blue) PCa samples. The y-axis represents sample beta values for each individual CpG probe, after covariate adjustment. The FDR significance in the difference of mean beta values between the two groups, as per a t-test, is shown.

Validating our ancestry-associated differential methylation findings, we performed equivalent analyses in a tumour-derived EPICv2 “ancestry-associated validation cohort” for 35 African (all southern African) and 22 non-African (3 European southern Africans, 1 Asian- and 18 European Australians) patients, while acknowledging significant confounding between ancestry and tumour purity, minimising appropriate adjustment. Validation cohort results can be found in Supplementary Figs. 8-11 and Supplementary Tables 10-13.

### African prostate tumour-associated differential methylation discovery

Using the African-specific EPICv2 prostate tissue-derived “tumour-associated discovery cohort”, including 93 southern Africans either with (*n* = 35) or without (*n* = 58) clinicopathologically confirmed PCa (Table 1), distinguished patients by DNA methylation profile through PCA visualization, with the spread in PC2 attributable to non-tumour stromal and immune cell content (Fig. 4a, Supplementary Figs. 12-13, Supplementary Table 14). Furthermore, we found prostate tumours to be associated with increased genome-wide methylation in CpG dense regions, while conversely associated with decreased genome-wide methylation at repetitive elements (LINE-1, LTR, Alu) (Fig. 4b, Supplementary Fig. 14), with the latter showing 27.50% collective overlap with common PMDs.

**Fig. 4:**
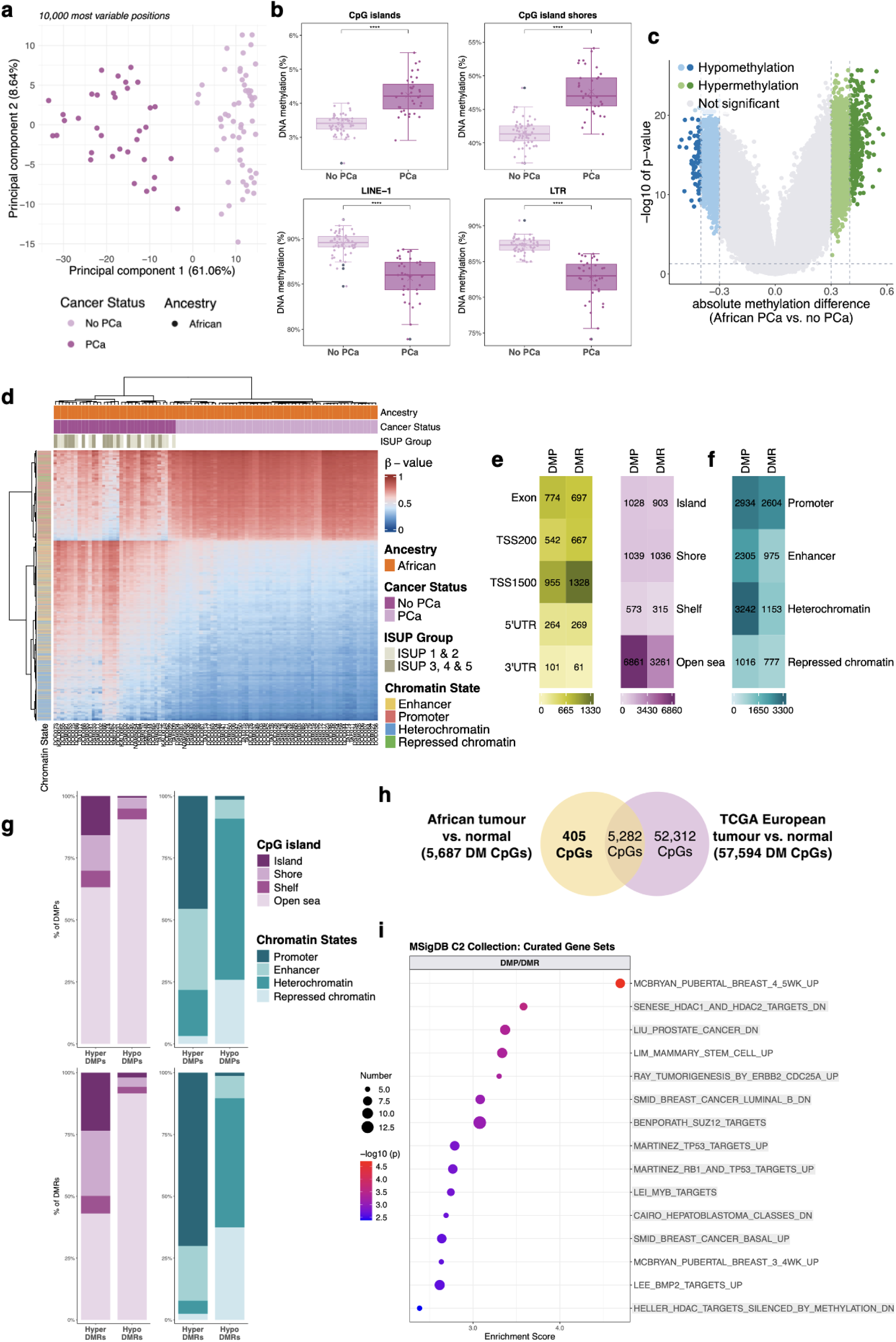
African-specific prostate tumour *versus* normal tissue differential methylation. **a** PCA for the 10,000 most variable positions for the EPICv2 African “tumour-associated discovery cohort” by cancer status: those with PCa (*n* = 35), and those without PCa (*n* = 58). **b** Boxplots of global DNA methylation levels in African prostate tumour *versus* normal tissue samples across CpG islands, CpG island shores, LINE-1 repetitive elements and LTRs. Each dot represents the median methylation value for each patient. The “X” indicates group mean. **c** Volcano plot representing 9,501 DMPs by cancer status (prostate tumour versus normal tissue) significance (BH FDR *p* <0.05, |Δβ| ≥30% and 40%, dashed lines), including hypermethylated (green) or hypomethylated (blue), compared with non-significant (grey) DMPs. **d** DMP cluster analysis heatmap by cancer status, ISUP grade group and chromatin state. Rows represent CpG sites (*n* = 9,501) and columns, patients (*n* = 93). The methylation level is represented by a β-value between 0 (completely unmethylated, blue) and 1 (fully methylated, red). **e** Heatmap showing gene region (green) and CpG island-related region (purple) enrichment for DMP- and DMR-related CpG sites across various contexts. **f** Heatmap showing chromatin state enrichment of DMP- and DMR-related CpG sites across various contexts. **g** Stacked bar graphs of the percent overlap of DMPs and DMRs with various CpG island-related (purple) and chromatin state (blue) contexts. **h** Venn diagram illustrating shared and distinct significant differentially methylated CpGs (BH FDR *p* <0.05, |Δβ| ≥10%) identified in the African tumour-associated cohorts and TCGA-PRAD European cohort. **i** MSigDB (C2) enrichment for genes collectively annotated to hypermethylated DMPs and DMRs not identified within TCGA-PRAD European cohort for the top 15 terms (minimum 5 genes), with cancer-related terms highlighted in grey. Point size reflects the number of genes associated with each gene set. Association strength is denoted by – log10(*p*-value). Abbreviations: *DM*, differentially methylated; *DMP*, differentially methylated position; *DMR*, differentially methylated region; *ISUP*, International Society of Urological Pathology; *LTRs*, long tandem repeats; *MSigDB*, Molecular Signatures Database; *PCA*, principal component analysis; *TCGA*, The Cancer Genome Atlas; *TSS*, transcription start site; *UTR*, untranslated region; *|Δβ|*, absolute difference in mean methylation; ****, *p* ≤0.0001.

Through differential methylation analysis, we identified 9,501 tumour-associated DMPs (BH FDR *p* <0.05, |Δβ| ≥30%) (Supplementary Table 15), with hypermethylation dominance (66.64%) in tumour positive tissue (Fig. 4c), and DMP cluster analysis showing a clear distinction between tumour and non-tumour samples, with bias towards tumour-associated hypermethylation (Fig. 4d). Reminiscent of our ancestry-associated findings, tumour-associated DMPs were primarily located in non-CGI regions (72.21%), followed by CGIs and shores (21.76%), while showing predominant enrichment in heterochromatin (34.12%, *p* = 4.33 × 10^−13^), followed by promoters (30.88%), including active non-prostate (12.16%, *p* <2.2 × 10^−16^) and prostate lineage-specific (2.21%, *p* <2.2 × 10^−16^), and bivalent poised (16.51%, *p* <2.2 × 10^−16^) promoters, and enhancers (24.26%, *p* = 2.51 × 10^−12^) (Fig. 4e-f). To assess for possible functional importance, we determined DMP location relative to various genomic contexts (Fig. 4g, Supplementary Fig. 15). Consistent with our global analysis, CGI regions exhibit greater gain of methylation over loss. However, most hypo- and hypermethylated DMPs are in non-CGI regions (90.47% and 63.07%, respectively). Relative to annotated chromatin states, hypermethylated DMPs are mainly located in promoters (45.59%, *p* <2.2 × 10^−16^), enhancers (32.59%, *p* = 2.02 × 10^−7^) and heterochromatin (18.64%, *p* <2.2 × 10^−16^), with hypomethylated DMPs found mostly in heterochromatin (65.05%, *p* <2.2 × 10^−16^) and repressed chromatin (25.80%, *p* <2.2 × 10^−16^), with significant enhancer enrichment (7.63%, *p* <2.2 × 10^−16^).

Furthermore, we identified 707 tumour-associated DMRs comprising 5,444 CpGs (FDR *p* <0.05, |Δβ| ≥20%) (Supplementary Tables 16-17). Compared to non-tumour samples, 423 DMRs are hypermethylated and 284 hypomethylated in the African tumours, with overall distribution in promoter regions (47.22%) (Fig. 4e-f). Consistent with global CpG methylation, hypermethylated DMRs are biased to CGI regions (49.80%), while hypomethylated DMRs predominate within open sea regions (91.54%) (Fig. 4g). Relative to chromatin states, 70% (*p* <2.2 × 10^−16^) of hypermethylated compared to 1.42% of hypomethylated DMRs are located at promoters, with most hypomethylated DMRs distributed in heterochromatin (52.24%, *p* <2.2 × 10^−16^) and repressed chromatin (37.45%, *p* <2.2 × 10^−16^). Enhancers were enriched for both hyper- and hypomethylation (22.05%, *p* = 1.12 × 10^−11^ and 8.90%, *p* <2.2 × 10^−16^, respectively). Interestingly, both hypomethylated DMPs and DMRs show negligible promoter distribution (1.51% and 1.42%, respectively).

To consider African-specific differential methylation, we first performed differential methylation analysis between The Cancer Genome Atlas (TCGA)-PRAD European tumour and normal samples (see Methods) (57,594 DMPs, BH FDR *p* <0.05, |Δβ| ≥10%, Supplementary Table 18), and then conducted functional enrichment analysis on 405 African tumour-associated differentially methylated CpGs, selected for their overlap with our tumour-associated validation cohort but exclusion from TCGA-PRAD cohort (Fig. 4h, Supplementary Table 19). Again, only annotated genes with a prostate tissue methylation/expression correlation were included. Hypermethylated DMPs/DMRs were enriched for cancer-related gene sets (MSigDB C2 collection) defined as downregulated in (metastatic) PCa, downregulated upon HDAC knockdown (Fig. 4i) and silenced by methylation in several cancer types (Supplementary Table 20). Hallmark gene sets included TNF-α signalling and cholesterol homeostasis. Conversely, hypomethylated DMPs/DMRs exhibited sparse gene set enrichment, unsurprising given 50% of the hypomethylated CpGs fall within a common PMD. Not all pathways were significant after controlling for a 5% FDR. Utilizing the GeneHancer ‘Double Elite’ list, we verified DMP and DMR overlap with 396 validated promoter/enhancer elements. Two or more sources of evidence suggest a high likelihood of interaction between these regions and 749 target genes collectively (Supplementary Table 21), with these genes showing enrichment in gene sets of known TF targets, including *FOXO4*, and numerous cancer-associated pathways (Supplementary Table 22).

Of the 405 CpGs found to be African-specifically differentially methylated in this study, 27 CpGs displayed a DMP/DMR |Δβ| ≥20%, annotated to 16 genes: *GJB5*, *CD1E*, *C2orf88*, *SLC19A3*, *PROM1*, *ARL9*, *MIR575*, *SLC12A9*, *LRRC4*, *TACC1*, *HOXC4*, *ADCY4*, *PYCARD*, *CX3CL1*, *FBXO17* and *KLF8*. While 11 genes, including *PYCARD*, have PCa associations (Supplementary Table 23), it appears that *GJB5*, *CD1E*, *SLC19A3*, *SLC12A9* and *FBXO17* have not been previously reported in relation to PCa in published studies. However, we found two CpGs to be of particular interest as potentially unknown African-specific PCa CpG targets: cg04742719 (*SLC12A9*) and cg11970458 (*PYCARD*) (Fig. 5). We also observed tumour-associated hypermethylation amongst several of the most extensively studied and independently validated DNA methylation biomarkers for PCa^7^, including *GSTP1*, *CCND2* and *PDLIM4*.

**Fig. 5:**
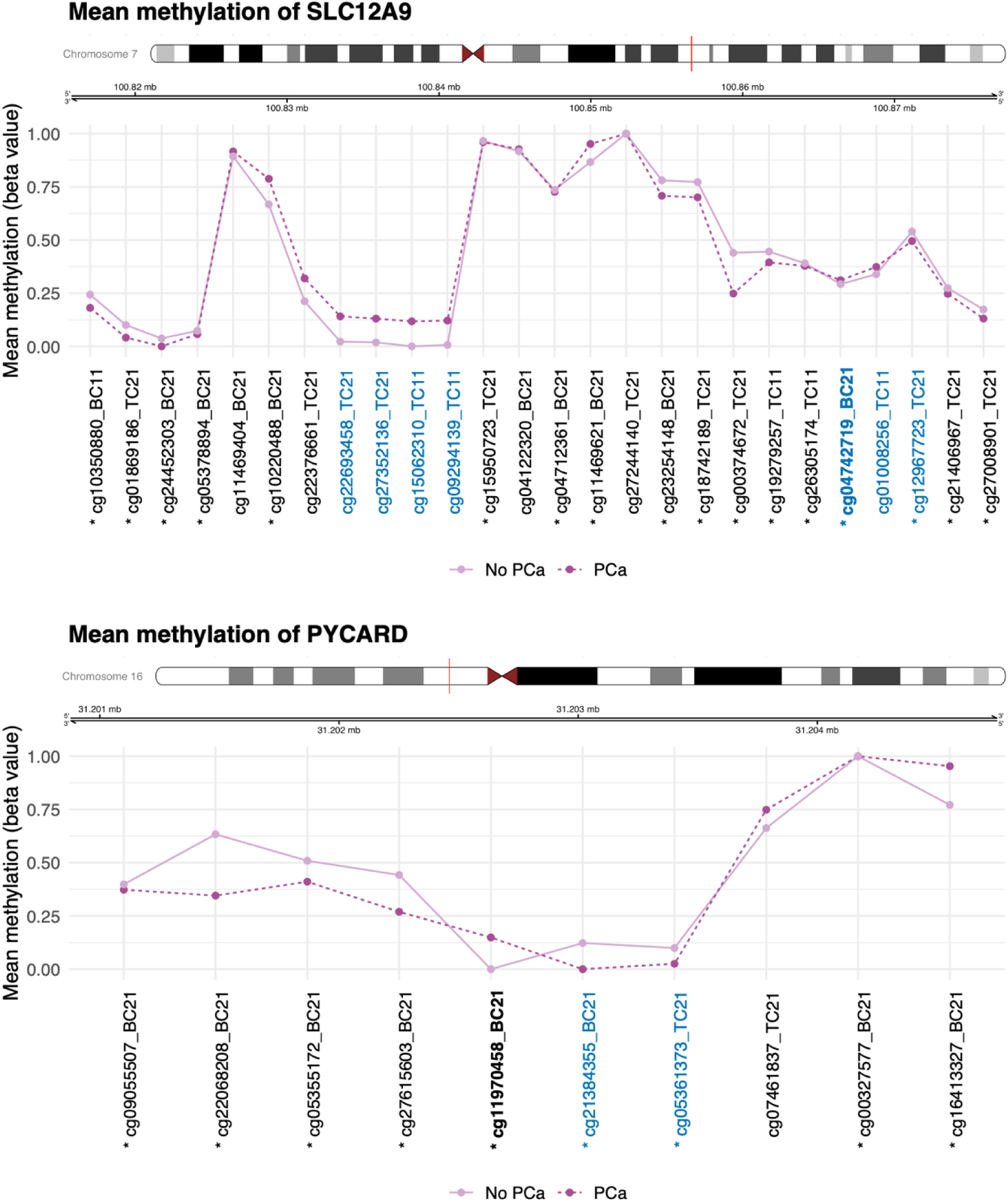
African prostate tumour *versus* normal tissue gene methylation plots highlighting two individual potentially unknown African-specific CpG prostate cancer targets. Differential methylation over noteworthy genes *SLC12A9* and *PYCARD*, highlighting (in bold) significant potentially unknown African-specific prostate tumour *versus* normal CpG targets. Probes overlapping CpG islands are shown in blue, the y-axis represents the mean beta value per group for each individual CpG probe, after covariate adjustment, and differential mean methylation significance between groups, determined by FDR *p* <0.05, is represented by an asterisk (*).

Equivalent analyses were performed using our independent “tumour-associated validation cohort” including EPICv1 prostate tissue-derived data from southern African men either with (*n* = 22) or without (*n* = 7) clinically confirmed PCa, again, acknowledging confounding in this cohort between our variable of interest and stromal cell content. Validation cohort results can be found in Supplementary Figs. 16-20 and Supplementary Tables 24-27.

## Discussion

The inclusion of African populations across the rich geographic diaspora is crucial for uncovering the genetic and environmental factors that contribute to variation in PCa risk and associated clinical adversity. This is further enhanced through the study of epigenetic modifications, with a focus on DNA methylation, a common feature of tumorigenesis and associated environmental exposures. However, studies focused on elucidating the role of differential methylation associated with African-associated PCa health disparities, has largely been restricted to the context of the United States^3^, with a notable lack of data for the African continent. Representing extreme genetic and environmental diversity^21^, as well as the greatest global PCa mortality rates^1^, here we focus on southern Africa. Establishing a regionally relevant filtering resource for Illumina EPIC-generated DNA methylation datasets, we provide compelling evidence for ancestry over geographic-associated prostate tumour methylation and extensive African-specific heterogeneity, while identifying African-relevant differentially methylated PCa targets.

Given the shared evolutionary history (Out of Africa) and in turn, genetic similarity between Asian and European individuals, both distant from Africans^30^, it appears genomic characterisation by ancestry, including in our study multi-generational European southern Africans, is mirrored in the PCa methylome. Overall, African-derived tumours displayed greater DMP and DMR hypermethylation, with DMRs biased towards CGIs and promoters. Epigenetic silencing of tumour suppressor genes through promoter CGI methylation is a hallmark of PCa severity, consistent with the typical inverse correlation between promoter CGI methylation and gene expression^31^. Greater hypermethylation of DMRs in protein-coding regions in Africans suggests that epigenetic silencing, while characteristic of PCa across ancestries, may occur more frequently in Africans, possibly contributing to more aggressive phenotypes. Consistent with hypermethylation-induced silencing, gene ontology analysis of hypermethylated DMP/DMR-associated genes showed enrichment for gene sets reported as downregulated in metastatic prostate tumours (“CHANDRAN_METASTASIS_DN”)^32^. Similarly, our tumour-associated hypermethylated DMRs were also mostly found within CGI-related regions and promoters, with overlapping associated gene set enrichment.

In contrast, ancestry-associated hypomethylated DMRs, often overlapping enhancers and promoters, may indicate epigenetically activated sites in African prostate tumours. This epigenetic activation, particularly of oncogenes, is another known tumorigenic event correlated with PCa severity^31^. In line with this, hypomethylated DMPs/DMRs were significantly enriched for gene sets involved in EMT, representing upregulated immune responses and target genes of oncogenic TFs including E2F5 and E2F2. Moreover, predominant enrichment of aberrant methylation in heterochromatin may point to disrupted chromatin structure maintenance and consequent genomic instability^33^. Transcriptionally inactive heterochromatin is hypoacetylated, catalysed by HDACs that remove acetyl groups from histone tails. Our ancestry- and tumour-associated hypermethylated DMPs/DMRs were enriched for gene sets downregulated in response to HDAC inhibition^34,35^ or possessing HDAC activity, suggesting diminished histone deacetylation activity in African tumours may contribute to genomic instability, another established feature of African PCa^36^, along with African-specific epigenetic machinery alteration of HDACs^37^.

Analogous to our ancestry associations, African-derived tumours over non-tumours were primarily hypermethylated, with DMPs biased to non-CGI regions and displaying substantial enhancer distribution. Enhancers regulate gene expression distally via interactions with TFs and transcriptional machinery, and DNA looping to achieve proximity with gene promoters. Cancer significantly alters enhancer methylation, affecting (cancer-associated) target gene expression^25^. We again observed target gene enrichment for “CHANDRAN_METASTASIS_DN”, indicating both direct gene dysregulation through aberrant methylation and indirect effects via epigenetically altered interacting enhancers.

While ancestry-associated enhancer target genes included targets of *ETS2* proto-oncogene and *FOXO4* tumour suppressor, validated tumour-associated hypermethylated promoter/enhancer elements included target genes involved in tumour suppressor binding motifs (FOXO4, SMAD2, TCF21, RB1, TP53). FOXO4 suppresses metastasis by inactivating the oncogenic PI3K/AKT pathway^38^. Given DNA methylation represses enhancer activity^39^, reduced tumour suppressor function in African PCa may occur through methylation of enhancer binding regions. This suggests an oncogenic mechanism whereby enhancer hypermethylation disrupts TF recruitment, binding dynamics and enhancer-promoter looping, ultimately altering target gene expression.

Conversely, tumour-associated targets of hypomethylated regulatory regions belong to upregulated cancer-associated gene sets, developmental pathways and oncoprotein binding motifs, including E12 (encoded by *TCF3*, a PCa oncogene). Exhibiting developmental pathway involvement, hypomethylated DMPs/DMRs were enriched in heterochromatin and repressed/inactive chromatin states. In differentiated cells, facultative heterochromatin typically represses developmental genes, dynamic to developmental and/or environmental cues^40^. We speculate that heterochromatin hypomethylation in African tumours creates permissive chromatin states that expose developmental genes to transcriptional activation. This effect, combined with epigenetic activation of enhancers targeting these genes, promotes pluripotency and carcinogenesis. This mechanism may predominate in African tumours, as hypomethylation occurred primarily in intergenic regions, suggesting oncogenic activation beyond promoters. Notably, enhancer target genes showed slightly greater enrichment for developmental pathways than DMPs/DMRs themselves, indicating enhancer (*versus* promoter) methylation status better reflects African-specific developmental gene dysregulation, while DMP/DMR-associated genes better indicate immune pathway alterations.

Top hit ancestry-associated DMPs/DMRs impact genes involved in cell adhesion and neuronal development (*NBEA*, *CHL1*), development and cell differentiation (*MAB21L1*, *RDH11*), immune signalling (*SYK*), protein degradation (*RNF217*) and angiogenesis inhibition (*TFPI2*). Hypermethylated in African tumours, *NBEA* has been shown to be downregulated in PCa^41^, *CHL1* implicated in PCa predisposition^42^ and *RNF217* has shown reduced copy number/expression in radioresistant LNCaP^43^.

Identifying unknown ancestry- or tumour-associated African-specific PCa target genes, these include functionality ranging from developmental signalling (*EVC2*, *CHSY1*), cell cycle regulation (*SPDYA*), ion homeostasis (*SLC12A9*), glucose metabolism (*GALM*), to inflammatory response and apoptosis (*PYCARD*). *CHSY1* shows gene body hypermethylation (positively correlated with expression) in African tumours and correlates with poor prognosis in gastric cancer^44^, while accumulation of its chondroitin sulfate chains is linked to prostate tumour aggressivity^45^, although no research has reported PCa gene-specific alterations. Hypomethylated in African tumours, *GALM* overexpression is associated with poor prognosis in glioma^46^. While *SPDYA* (also *Spy1*) typically shows overexpression in ovarian cancer^47^, its hypermethylation in African tumours suggests overexpression may be a mechanism unique to non-African tumorigenesis. Another notable observation includes *EVC2*’s involvement in the cancer-implicated Hedgehog signalling pathway^48^, though its specific role in PCa requires further investigation. While *PYCARD* (cg11970458), but not *SLC12A9* (cg04742719), exhibits a role in PCa, the observed tumour *versus* normal hypermethylation of these particular CpGs appears to be African-specific. Upregulated solute carrier *SLC12A9* is associated with aggressive uveal melanoma^49^, while *PYCARD* encodes an apoptosis-inducing factor and is characteristically suppressed by hypermethylation in aggressive PCa^50^. Notably, no ancestry-related differentially methylated genes typically reported in African American PCa overlapped with our southern African cohort, likely reflecting ancestral genetic divergence^21^, reiterating the need for further inclusion across the broad African diaspora.

Our study highlights challenges associated with DNA methylation studies across Africa. Observing substantially more African *versus* European variant^22^ overlap with array probes, although notably reduced for the updated EPICv2, raises the need for population-tailored approaches across Sub-Saharan Africa to minimise false positives due to genetic confounding. Further studies should explore applicability in larger, unrelated cohorts, for concordance between pan-African genomic variation^51^. While our African ancestral prostate tumours showed extensive between tumour DNA methylation heterogeneity, seven of 72 (9.72%) non-cancer pathology-defined biopsies resembled tumour-specific methylation. Assuming misclassification, it is notable that African American men are more likely to present with an anterior presenting tumour^52^, which is harder to detect using digitally guided transrectal prostate biopsy, the standard diagnostic approach for our southern African patients. Besides misclassification, this may also result in reduced tumour purity, possibly accounting for the heightened cell type heterogeneity observed. We also acknowledge that while EPIC arrays have substantial overlap, differences in probe coverage limit array version validation to shared probes, potentially missing significant findings unique to each. Our analysis is further limited by African-specific CpG target identification using available TCGA 450k array European data, losing valuable information, while lack of patient-matched RNA and ChiP-seq data, limits for direct correlation between methylation changes and gene expression. The latter would be beneficial to confirm tissue-specific enhancer and chromatin state (in)activation. Finally, the absence of matched tumour genomic data, such as *TMPRSS2-ERG* fusion status, although reportedly less frequent in southern African-derived tumours^53^, precluded relevant adjustments.

In conclusion, we found African prostate tumours to exhibit distinct methylation patterns akin to aggressive disease, characterized by three key features (Fig. 6): (i) widespread gain of methylation with targeted silencing of PCa-relevant metastasis-related and tumour suppressor genes; (ii) aberrant methylation at distal enhancers disrupting TF interactions, with bias towards metastasis- and tumour suppressor binding motifs; and (iii) global hypomethylation of heterochromatic regions promoting developmental gene activation (aided by enhancer activation), influencing cell identity. Similarities between our ancestry- and tumour-associated analyses reflect tumorigenesis as the driver for African-specific methylation. The concurrent distribution of hyper- and hypomethylation at regulatory elements, coupled with activation of typically repressed chromatin, underscores the extensive regulatory plasticity of these intergenic regions driving genomic instability in African tumorigenesis. Substantial loss of methylation within repressed chromatin states emphasizes the importance of analysing regions distal to promoters, while highlighting how African-specific tumorigenesis emerges from complex genome-wide interactions rather than isolated gene dysregulation. As such, we demonstrate how African diversity holds rich potential to uncover unknown insights.

**Fig. 6:**
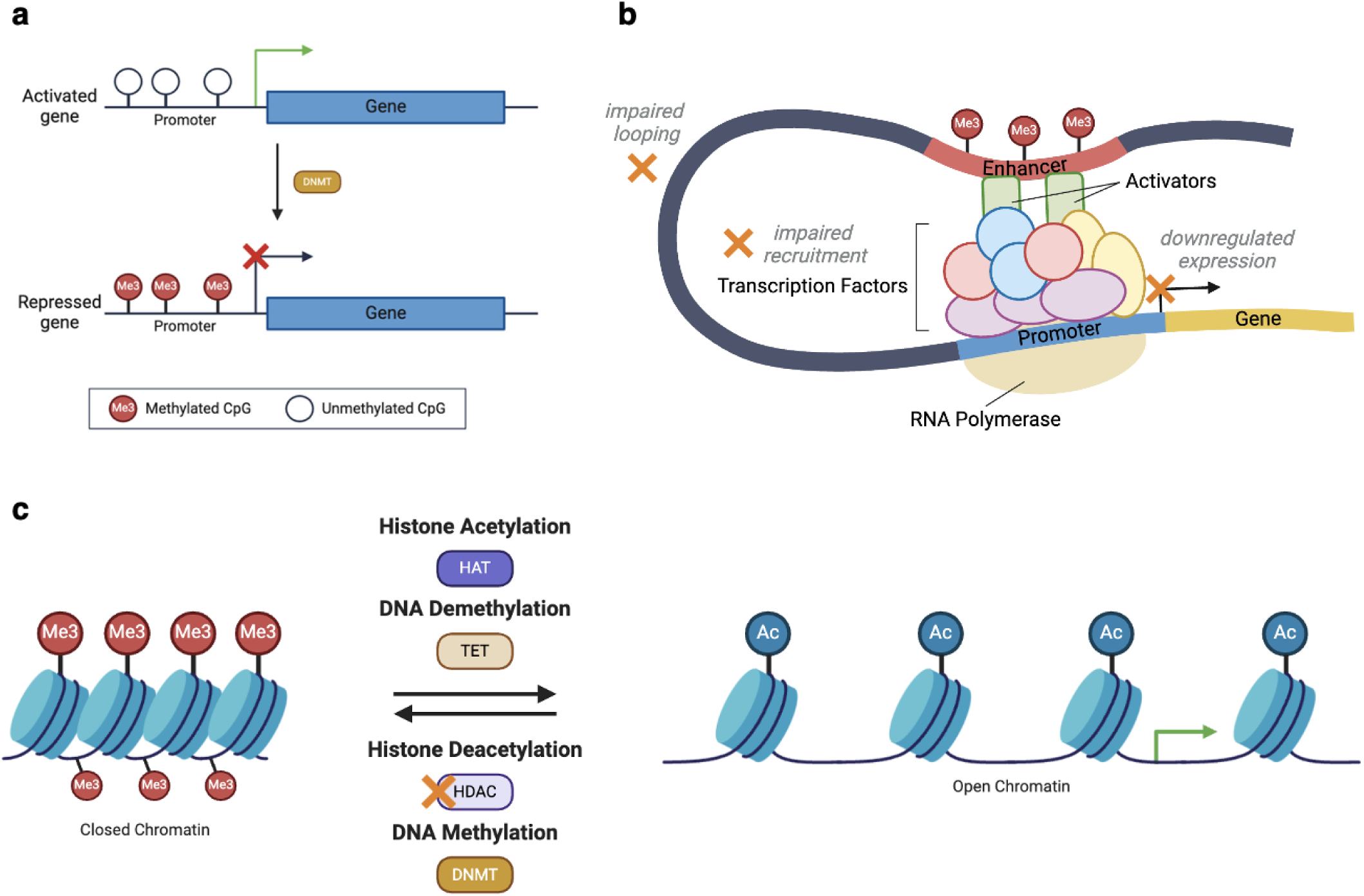
Overview of DNA methylation-driven epigenetic aberrations in African tumours. **a** Promoter hypermethylation-induced epigenetic silencing of (tumour-suppressor) genes. **b** Enhancer hypermethylation disrupts transcription factor (e.g. FOXO4) recruitment and binding dynamics, and impairs enhancer-promoter looping, ultimately altering target gene expression. Target genes of aberrantly methylated enhancers were often involved in tumour suppressor binding motifs. **c** DNA hypomethylation of closed chromatin promotes a more permissive chromatin state, which may be aided by diminished HDAC activity, collectively contributing to genomic instability. Abbreviations: *DNMT*, DNA methyltransferase; *HAT*: histone acetyltransferase; *HDAC*: histone deacetylase; *TET*: ten-eleven translocation.

## Methods

### Participant recruitment, sample processing and ethics

Written informed consent was obtained from 349 study participants recruited as part of the Southern African Prostate Cancer Study (SAPCS, *n* = 308), with approval granted by the University of Pretoria Faculty of Health Sciences Human Research Ethics Committee (HREC), with US Federalwide Assurance (FWA00002567 and IRB00002235 IORG0001762), in South Africa (HREC 43/2010) or the Ministry of Health and Social Services (MoHSS) in Namibia (17/3/3HEAF003). Additional Institutional Review Board (IRB) approval for SAPCS study recruitment was granted by the Human Research Protection Office (HRPO) of the US Army Medical Research and Development Command (E02371.2a TARGET Africa and E03333.1a and E05986.1a HEROIC PCaPH Africa1K in South Africa, and E03333.1b and E05986.1b HEROIC PCaPH Africa1K in Namibia). In Australia, St Vincent’s Garvan Prostate Cancer Study participant recruitment (*n* = 41) was approved by St. Vincent’s Hospital HREC (SVH/12/231). Recruitment occurred either at diagnosis (southern African) or surgery (Australian), with patients clinicopathologically classified using International Society of Urological Pathology (ISUP) grade grouping either with or without PCa (Table 1). All patients were treatment naïve at time of biopsy sampling.

Snap-frozen prostate cores (and patient-matched whole blood) were shipped on dry-ice to the University of Sydney in Australia under a Republic of South Africa Department of Health Export Permit (National Health Act 2003; J1/2/4/2) and in accordance with institutional Material Transfer Agreement (MTA) and associated Collaborative Research Agreement (CRA). Irrespective of country of recruitment, all prostate tissue samples (245 SAPCS, 41 Australian) were processed and Qiagen-based DNA extracted at the University of Sydney (Hayes Lab), minimising for technical artifacts. Tissue-associated Illumina Infinium HumanMethylationEPIC array data (Illumina, CA, USA) was generated at the Australian Genome Research Facility (AGRF, 2.8%) or the University of New South Wales’ Ramaciotti Centre for Genomics (Australia, 97.2%). Data generation and epigenomic/genomic interrogation was approved by the St. Vincent’s Hospital HREC (SVH/15/227), the University of Pretoria Faculty of Health Sciences HREC (504/2022) and by the HRPO of the US Army Medical Research and Development Command (E02371 TARGET Africa; E03280.1a and E05984.1a HEROIC PCaPH Africa1K). Samples were excluded if they did not meet quality control criteria during DNA methylation data processing or due to low African tumour purity estimations, excluding for 32.5% of samples (93/286, all SAPCS), leaving 193 (152 SAPCS, 41 Australian) for further analysis.

### Genomic resources and genetic ancestry fraction determination

As described previously^5^, blood-derived patient-matched whole genome germline small variant (SNVs and indels, MAF >1%) data called for the 99 southern Africans (36 overlapping with 193 methylation inclusive southern Africans) were made available via the SAPCS Data Access Committee (DAC) for generating a southern African EPIC-array SNP-filtering reference resource. To determine genetic ancestry for the 193 methylation inclusive participants, 132,245 filtered and linkage disequilibrium (LD) pruned germline SNPs (MAF >0.05, Hardy-Weinberg Equilibrium 1e-6 cutoff, LD pruned for window size of 50, step size of 10, r² threshold of 0.1), were made available to determine participant ancestry fractions (see Supplementary Table 28). Additional reference population data was downloaded from the Human Genome Diversity Project (HGDP) and 1000 Genome Project (1KGP) subset of gnomAD v3.12^24^, including 20 CEU Europeans, 20 CHB Chinese, and non-matched patients from the Southern African Prostate Cancer Study^5^ (40 SAPCS). Using unsupervised ADMIXTURE v1.3.0^54^ analysis (*K* = 3 replicated in 10 runs with average 0.524 cross-validation (CV) error), patient ancestries were assigned as “African” (*n* = 119, 2 Namibian, 117 South African) with >93% contribution (allowing for one Namibian outlier with 16% European fraction); “European” (*n* = 48, 18 Australian, 2 Namibian, 28 South African) if >90% contribution (allowing for a single outlier with an 11.1% Asian fraction) and “Asian” (*n* = 23, all Australian with >99% Asian fraction); with a single African-Asian admixed individual (*n* = 1 South African). A single African sample was excluded from estimations due to low coverage. Genetic ancestry for the 99 southern African reference participants (36 patient overlap) was determined using the same SNP set and reference data (Supplementary Table 29) and was replicated in 8/10 runs at *K* = 3, with a mean CV of 0.549. All patients were assigned “African” ancestry, with a majority (*n* = 96, 86 South African, 10 Namibian) with >87% African contribution, and three Namibian samples with between 14.9 to 22.7% European contribution. Namibians and South Africans were merged geographically for analyses as “southern African”.

### Establishing southern African SNP-filtering resources

Our workflow for identifying probe overlap with African variants is shown in Supplementary Fig. 1. To produce the EPICv1 filtering resource, 99 individual African germline VCF files were lifted over from hg38 to hg19 genome build (Picard v2.21.9 LiftoverVcf^55^). Individual VCF files were then merged into a single consensus VCF file, filtered for only those variants that pass quality control filters (bcftools^56^ v1.9), and chromosome notation style converted to that of NCBI. The VariantAnnotation and GenomicRanges packages^57,58^ were used to parse the single merged EPICv1 VCF file and extract all SNP and indel variants overlapping EPICv1 probes. As per Zhou *et al.*’s^19^ recommendations, we examined variants according to their probe position: (i) variants overlapping target CpG sites; (ii) variants overlapping single base extension (SBE) sites for Infinium type I probes; and (iii) variants overlapping the probe body within 5 bp from their targets. Results were filtered to include genetic variants with a maximum minor allele frequency > 0.01^19^ (see Supplementary Data 1-3). Barring lift over, the EPICv2 African filtering resource was generated using the same workflow and referencing the appropriate (hg38) Illumina manifest, excluding “chr0” and “chrM” probes (see Supplementary Data 4-6).

### Determining African-relevant CpG probe confounding

To determine the genomic distribution of African variant-confounding probe locations relative to specific features, EPICv1 probes were annotated using the Illumina manifest file “MethylationEPIC_v-1-0_B4.csv” mapped to hg19, and EPICv2 probes, using the “EPIC-8v2-0_A1.csv” manifest file mapped to hg38. Where EPICv2 probe annotation was unknown, we used Gencode data Release 25 (GRCh38.p7), downloaded in December 2024 from https://ftp.ebi.ac.uk/pub/databases/gencode/Gencode_human/release_25/gencode.v25.annotation.gtf.gz, to extract gene body and intergenic regions using the GenomicRanges package. We defined intergenic regions as regions mapping outside known genes. Enhancer elements were downloaded in December 2024 from the FANTOM5 enhancer atlas: https://fantom.gsc.riken.jp/5/datafiles/latest/extra/Enhancers/ (hg19) and https://fantom.gsc.riken.jp/5/datafiles/reprocessed/hg38_latest/extra/enhancer/ (hg38), comprising 65,423 and 63,285 enhancers from 1,829 human tissue samples, respectively^59^.

### Determining array version confounding

Matched African EPICv1 and EPICv2 replicate pairs were merged using RnBeads (v2.23.0)^60^, identifying shared probes across platforms by genome coordinate position. After merging raw datasets, correlation between 721,435 CpG sites common to both arrays was assessed for each replicate pair. For interrogation of array version as a confounder, raw EPICv1 (*n* = 78) and EPICv2 (*n* = 115) datasets were normalized using the “scaling.internal” method, background subtraction performed with the “sesame.noobsb” method, and datasets merged thereafter.

### Establishing discovery and validation methylation cohorts

Avoiding for array version confounding, tissue-derived HumanMethylationEPIC BeadChip v1.0 or v2.0 were analysed separately, allowing the larger resource to form the discovery arm and the smaller for validation. Consequently, the ancestry-associated discovery cohort consists of 70 EPICv1 samples and the validation cohort 57 EPICv2 samples. Conversely, the tumour-associated discovery cohort is made up of 93 EPICv2 samples and the validation cohort comprises 29 EPICv1 samples (Table 1). Only discovery cohort observations that were consistent with the validation cohort were presented.

### Array version-specific data processing, quality control and analyses

Using RnBeads and relevant annotation packages (hg19 for EPICv1 or hg38 for EPICv2, both v1.37.0), data was normalized using the “scaling.internal” method, followed by “sesame.noobsb” background subtraction. Samples were filtered for probes with detection *p*-values >0.05, probes with a bead count <3, default RnBeads SNP-overlapping probes, probes not in a CpG context and probes with missing values in >1% of samples. Cross-reactive probe filtering was performed manually as per Pidsley *et al.*^22^ for EPICv1, and Peters *et al.*^23^ for EPICv2. Probes located on sex chromosomes were retained. Samples were considered low quality if they displayed a missing probe fraction >0.2%.

Probes and samples not meeting quality control criteria were removed from subsequent analyses, with 717,828 probes and 109 samples in the EPICv1 dataset, and 794,018 probes and 173 samples in the EPICv2 dataset remaining for analyses. Filtering summaries are available in Supplementary Table 30. Normalized β-values were used for all subsequent statistical analyses.

### Cellular deconvolution of methylation data

To account for cell type heterogeneity, using the Houseman algorithm^61^ in RnBeads for reference-based deconvolution, we estimated cell type composition in bulk tumour samples, using DNA methylation profiles from sorted cell types as a reference^62^. Cell type proportion estimates for 193 samples can be found in Supplementary Table 28. To limit compounded heterogeneity introduced by the genomically-diverse African samples, only African tumour samples belonging to the highest tumour purity quartile were retained for further analyses (*n* = 57).

### Repetitive element, partially methylated domain (PMD) and chromatin state annotation of CpG sites

Annotation data for repetitive elements (Alu, LINE-1 and LTR) corresponding to the RepeatMasker database, available through AnnotationHub^63^, were downloaded using the REMP package (v.1.30.0)^64^. Annotations for common PMDs were retrieved from ref.^65^.

To overlay the regulatory context of CpG sites, we utilized a prostate tumour-derived chromatin state annotation of the human genome, as per ref.^26^. For simplification where relevant, “Active non-prostate lineage enhancer”, “Active prostate lineage-specific enhancer”, “Bivalent poised enhancer”, “Primed non-prostate lineage-specific enhancer” and “Primed prostate lineage enhancer” were collectively classed as “enhancers”; and “Active non-prostate lineage promoter”, “Active prostate lineage-specific promoter” and “Bivalent poised promoter”, as “promoters”.

### Identifying differentially methylated positions

For each cohort comparison, DMPs were identified by performing linear modelling with the limma package (v3.61.9)^66^, while adjusting for covariates where possible. For ancestry-associated analyses, age and tumour purity were adjusted for in the discovery cohort, and age in the validation cohort. In the tumour-associated analyses, age, stromal and immune cell content were adjusted for, while age and immune cell content were adjusted for in the validation cohort. Multiple testing correction was performed with the Benjamini–Hochberg (BH) false discovery rate (FDR) method, with probes deemed significantly differentially methylated if they displayed an FDR cutoff of *p* <0.05 and an absolute Δβ (i.e. absolute difference in mean methylation between two groups) ≥20% (ancestry-associated analysis) or ≥30% (tumour-associated analysis). Hierarchical clustering analysis was performed on significant DMPs using Euclidean distance and ward.D2 linkage.

### Identifying differentially methylated regions

DMRs were identified for each cohort comparison using matched regression models to those implemented in the respective DMP analyses. The package DMRcate (v3.1.9)^27^ was used to identify DMRs using a bandwidth smoothing window of 1000 bp, a bandwidth scaling factor of 2 and a minimum of five consecutive CpGs to define DMRs. DMRs with a minimum smoothed FDR cutoff of *p* <0.05 and an absolute Δβ of ≥10% (ancestry-associated analysis) or ≥20% (tumour-associated analysis) were considered significant.

### Enrichment analysis

Using the gsameth and gsaregion functions of the missMethyl package (v1.39.14)^67^, we identified Molecular Signatures Database (MSigDB, v.7.1)^28^ gene sets that were significantly enriched with methylation changes identified during DMP and DMR (both minimum |Δβ| ≥10%) identification. MSigDB gene sets were downloaded in November 2024 from https://bioinf.wehi.edu.au/MSigDB/v7.1/. Using a subset of prostate (tumour)-derived TCGA-PRAD samples (*n* = 309), patient-matched methylation (Illumina HumanMethylation450) and gene expression (STAR - Counts) quantification data were downloaded in January 2025 using the TCGAbiolinks package’s (v2.34.0)^68^ GDCquery function and used to compute

Pearson correlations between CpG beta and log2 expression values. Patient-matched TCGA-PRAD samples were included if treatment-naïve, primary prostate gland-derived adenocarcinomas (*n* = 274), or normal solid tissue (*n* = 35). To determine the regulatory impact of significant DMPs/DMRs, only related genes with a significant correlation (FDR *p* <0.05, |*r* | >0.25) or not covered on the 450k array, were included for enrichment analysis. For our tumour-associated enrichment analysis, we further subset TCGA-PRAD methylation data to only include individuals of “white” (European) race (*n* = 249, 217 primary solid tumours, 32 solid tissue normal), performed tumour *versus* normal differential methylation analysis using *limma*, while adjusting for age at diagnosis, and used resultant European-related DMPs (FDR *p* <0.05, |Δβ| ≥10%) to identify uniquely African tumour-associated differentially methylated sites limited to those covered on the 450k array, then used as input. The missMethyl package adjusts for the underlying distribution of probes on the array, with genes annotated to DMP and DMR probes compared to lists of genes in each curated MSigDB gene set to identify those that are statistically over-represented.

DMPs and DMRs were flagged for overlap with GeneHancer “Double Elite”^29^ regions i.e. both GeneHancer elements and genes having associations derived from at least two information sources. Putative target genes for DMP- and DMR-overlapping enhancers were tested for gene set enrichment in MSigDB gene sets using the RITAN and RITANdata packages^69^. Using TCGA-PRAD expression data, detailed above, target genes were filtered for those considered expressed in the prostate, defined by a mean log2 expression value ≥2 per gene. We defined the background as all GeneHancer “Double Elite” genes and considered terms with an FDR *p* <0.05 as significant.

### Statistical analysis

All statistical analyses were performed using R software (v4.4.1). Cross-platform methylation correlation was calculated using Pearson’s correlation coefficient. Probes with an FDR <0.05 were considered significantly differentially methylated between platforms. Relative to non-significant differentially methylated sites, significant DMP/DMR enrichment across various genomic contexts was calculated using Fisher’s exact test. For repetitive element and CGI-related region boxplots, group means were compared using a t-test.

## Data availability

Methylation data files generated in this study have been deposited in the Gene Expression Omnibus (GEO) under accession number GSEXXXXXX. Ancestry informative variant data has been uploaded to the European Variation Archive (www.ebi.ac.uk/eva) under accession code GCSTXXXX and the SNP-filtering reference data under access code GCSTXXXX, source data provided. All other data supporting the key findings of this study are available within the article and Supplementary Information.

## Code availability

R scripts for parsing and identifying EPIC probes that overlap southern African variants have been deposited at GitHub and can be accessed at https://github.com/j-craddock/African-variants-overlapping-EPIC-probes.

## Acknowledgements

We are forever grateful to the study participants for their contribution to this research, as well as both current and past clinical support staff who have contributed to the Southern African Prostate Cancer Study (SAPCS), Namibian-SAPCS and the Garvan Institute St Vincent’s Prostate Cancer Biobank. We further acknowledge the SAPCS and Garvan-St Vincent’s Biobank Managers Tumisang Mbeki (University of Pretoria, South Africa) and Anne-Maree Haynes (Garvan Institute of medical Research, Australia), respectively, Dr Clare Stirzaker and Dr Elena Zotenko (Garvan Institute of medical Research, Australia) for sharing their original code for the identification of EPIC probes overlapping genetic variants, Jue Jiang (Ancestry & Health genomics laboratory, University of Sydney, Australia) for providing germline variant calling, and Dr Matthew Freedman and Dr Xintao Qiu (Dana–Farber Cancer Institute, Harvard Medical School, USA) for sharing the prostate cancer ChromHMM annotation. This study was supported by funding received from the U.S.A. Congressionally Directed Medical Research Programs (CDMRP) Department of Defense (DoD) Prostate Cancer Research Program (PCRP) Idea Development Award (PC200390, TARGET Africa to V.M.H.) and a HEROIC Consortium Award (PC210168 and PC230673, HEROIC PCaPH Africa1K to V.M.H. and M.S.R.B.), a U.S.A. National Institute of Health (NIH) National Cancer Institute (NCI) Award (1R01CA285772-01 to V.M.H.); U.S.A. Prostate Cancer Foundation (PCF) Challenge Award (2023CHAL4150 to V.M.H.) and National Health and Medical Research Council (NHMRC) Ideas Grants (APP2001098 to V.M.H. and M.S.R.B.; APP2010551 to V.M.H.). J.C. was further supported by the National Research Foundation of South Africa (PMDS22070633683) and V.M.H. by the Petre Foundation via the University of Sydney Foundation, Australia. This study will form part of a PhD dissertation of J.C..

We further acknowledge **HEROIC PCaPH Africa1K** co-principal investigators: Gail S. Prins^1^ and Peter Mungai Ngugi^2^; developmental team leads: Weerachai Jaratlerdsiri^3,4^, Winstar Mokua Ombuki^2^, Daniel M. Moreira^1^ and Ikenna C. Madueke^1^; SAPCS Resource Working Group members: Maphuti Tebogo Lebelo^5^, Tumisang M. N. Mbeke^5^, Muvhulawa Obida^5,6^, Martin Obida^6^, Raymond Campbell^7^, Mulalo B. Radzuma^8^, Golda Stellmacher^9^, Jessie Gamxamub^9^, Reginold M. J. Menoe^10^ and Jeff John^11^; Genomics and Data Science Working Group members: Jue Jiang^3^, Tingting Gong^3^, Korawich Uthayopas^3^, Kazzem Gheybi^3^, Ruotian Huang^3^, Kangping Zhou^3^, Umuna Maendo^3,12^, Massimo Loda^13^, G. Nicolo’ Fanelli^13^, David C. Wedge^14^, Avraam Tapinos^14^, Vivien Holmes^14^, Robert G. Bristow^14^, Daniel S. Brewer^15,16^, Abraham Gihawi^15^, Colin S. Cooper^15,17^, Rosalind A. Eeles^17^ and Zsofia Kote-Jarai^17^; and other key Consortia members (alphabetical order): Maria Argos^18^, Irene Barnhoorn^19^, Lynn Birch^1^, Muriuki Elias Nyaga^20^, Micah O. Oyaro^2^, Joyce Shirinde^5^, Douglas I. Walker^21^, Edwin O. O. Walong^22^, Githui Shila Wanjiku^2^, and Margaret Quaid^18^.

## HEROIC PCaPH Africa1K Associated Author Affiliations

^1^Department of Urology, University of Illinois at Chicago, Chicago, IL, USA; ^2^East Africa Kidney Institute, Department of Urology, University of Nairobi, Nairobi, Kenya; ^3^Ancestry and Health Genomics Laboratory, Charles Perkins Centre, School of Medical Sciences, Faculty of Medicine and Health, University of Sydney, Camperdown, NSW 2050, Australia; ^4^Computational Genomics Group, Charles Perkins Centre, School of Medical Sciences, Faculty of Medicine and Health, University of Sydney, Camperdown, NSW 2050, Australia; ^5^School of Health Systems and Public Health, University of Pretoria, Pretoria, South Africa; ^6^Tshilidzini Hospital, Shayandima, Thohoyandou, Limpopo, South Africa; ^7^Department of Urology, University of Pretoria, Pretoria, South Africa; ^8^Department of Urology, Sefako Makgatho Health Science University, Dr George Mukhari Academic Hospital, Ga-Rankuwa, Gauteng, South Africa; ^9^Department of Urology, Windhoek Central Hospital, Windhoek, Namibia; ^10^Life Peglerae Hospital, Rustenberg, North West, South Africa; ^11^Frere Hospital, East London, South Africa; ^12^Botswana International University of Science and Technology, Palapye, Botswana; ^13^Department of Pathology and Laboratory Medicine, Weil Cornell Medicine, New York Presbyterian-Weill Cornell Campus, New York, NY, USA; ^14^Manchester Cancer Research Centre, University of Manchester, Manchester, UK; ^15^Norwich Medical School, University of East Anglia, Norwich, UK; ^16^Earlham Institute, Norwich, UK; ^17^The Institute of Cancer Research, London, UK; ^18^Department of Environmental Health, Boston University School of Public Health, Boston, MA, USA; ^19^University of Venda, Limpopo, South Africa; ^20^Meru County Referral Hospital, Moi University, Meru County, Kenya; ^21^Gangarosa Department of Environmental Health, Emory University, Atlanta, Georgia, USA. ^22^Department of Pathology, University of Nairobi, Nairobi, Kenya.

## Author Contribution Statement

Conceptualization: V.M.H.; Methodology and software: J.C., P.L., P.X.Y.S., C.G. and V.M.H.; Formal analyses and investigation: J.C.; Participant recruitment, clinical data and resources: S.B.A.M., P.D.S., H.E.A.F. and M.S.R.B.; Sample processing: M.M.H.; Data curation: J.C., M.M.H., and V.M.H.; Writing—original draft preparation and figures: J.C.; Writing revisions: J.C., P.L., P.X.Y.S., C.G. and V.M.H.; Study supervision: P.L., C.G. and V.M.H.; Project administration: S.M.P., M.S.R.B. and V.M.H.; Funding acquisition: M.S.R.B. and V.M.H. All authors have read and agreed to the published version of the manuscript.

## Competing interests

Hayes is a Member of Active Surveillance Movember Committee and received an honorarium from The Korean Urological Oncology Society for 2024 Annual Conference as a guest speaker.

